# New distributional data for the northern forestfly, *Lednia borealis* Baumann and Kondratieff, 2010 (Plecoptera: Nemouridae), Washington, USA

**DOI:** 10.1101/2021.01.25.428104

**Authors:** Candace E. Fallon, Emilie Blevins, Michele Blackburn, Taylor B. Cotten, Derek W. Stinson

## Abstract

The northern forestfly, *Lednia borealis* (Plecoptera: Nemouridae) is a rare montane stonefly believed to be endemic to Washington. The species, first recognized as a valid taxon in 2010, is the only member of the genus *Lednia* known from the state. Like other species in its genus, it is found in mid- to high-elevation cold water habitat, including lakes, glacial-fed streams, and rheocrenes (channelized springs). *Lednia* species in general appear to be rare or at least rarely collected. Because of their reliance on alpine and subalpine habitat, *Lednia* may be especially vulnerable to threats associated with climate change. However, relatively little is known about this species, and distribution data are scarce. From 2015 to 2019, 94 sites were surveyed in order to document unmapped populations of *Lednia borealis* to improve range and distributional information from montane areas of Washington State. In this paper, we share locations of *L. borealis* documented to date, including collections from eight newly documented *Lednia* sites in the Mt. Baker and Glacier Peak Wildernesses in the Cascade Mountains of Washington, and report recent COI barcoding results. We also provide updated details on the species’ distribution, highlight a confirmed habitat association with glacial edge meltwater, and provide recommendations for future surveys.

## INTRODUCTION

Stoneflies (Order: Plecoptera) are integral components of freshwater ecosystems, playing vital roles in nutrient cycling and decomposition and serving as food sources for other species such as fish and other larger insectivores (DeWalt and Ower 2019). Larvae are aquatic, living in well-oxygenated lakes, streams, springs, and seeps and feeding on submerged leaves, benthic algae, detritus, or other invertebrates. Larvae molt numerous times, typically undergoing 10 to 24 instars before emerging from streams as adults (Stewart 2009). Adults of almost all species are terrestrial and capable of flight. The larval stage can last from several months up to four years, whereas adults tend to live only a few weeks (Stewart 2009). Although some do not feed as adults, others may feed on algae and lichen attached to the bark of trees, or leaves, buds, and pollen grains from riparian vegetation (Stewart 2009, DeWalt and Kondratieff 2019). Little is known about the distribution, life history, and population status of many species.

Stoneflies are generally poorly studied, and this is particularly true for alpine-specialist species, whose habitats may be difficult to access and are often limited to cold, clear water in montane streams, springs, and seeps, many of which have glacial origins or influences. These species are rarely encountered or targeted in survey efforts, despite their potentially greater need for conservation attention resulting from their reliance on specialized and imperiled habitats, limited dispersal abilities, and endemism. Indeed, others have highlighted the limited ranges, general lack of knowledge, and overall conservation concern for alpine insect species (e.g., WDFW 2015, Hotaling et al. 2017, Birrell et al. 2020). Few baseline surveys have been conducted to determine distribution and general habitat requirements. Even with targeted searches, some of these species are difficult to detect and observations remain scarce. Short adult flight periods, condition dependent life cycles, local snow conditions, patchy distributions, and cryptic life histories all serve to complicate surveys (e.g., see Fallon and Blevins 2015, Fallon et al. 2017). Lednian stoneflies (*Lednia* spp.) are prime examples of these poorly understood, rarely encountered montane species. Until 2010, *Lednia* was considered a monotypic genus. However, increased research focus on the genus has resulted in descriptions of four high-elevation species from the western United States and Canada: *L. tumana*, *L. sierra*, *L. tetonica,* and *L. borealis* (Baumann and Kondratieff 2010, Baumann and Call 2012).

*Lednia* specimens collected in Washington were initially thought to represent a range expansion of *L. tumana*; however, further taxonomic work revealed Washington-collected specimens to represent a new species, *L. borealis* (Baumann and Kondratieff 2010), later confirmed as a sister clade to the *L. tumana/L. tetonica* clade (Hotaling et al. 2019). Similar to other species in the genus, *L. borealis* is known only from a small number of sites. As such, it was believed to be a Washington State endemic, although Baumann and Kondratieff (2010) noted that it may occur in the southern Cascade Range in Oregon and northern California. However, these areas have not been well surveyed for high elevation stoneflies. A single adult female specimen collected from O’Brien in southwestern Oregon has been identified to species as *Lednia borealis* based on geographical locality (CNC 2020), although the location contradicts all other known habitat associations for the species and is highly disjunct, suggesting the need for further investigation. To the north, recent evidence indicates that *Lednia borealis* also occurs into central British Columbia (Green et al., in preparation). *Lednia borealis* is considered critically imperiled both nationally and at the state level in Washington (NatureServe 2020). It is also included as a Species of Greatest Conservation Need in the Washington State Wildlife Action Plan (WDFW 2015). It has been collected primarily from high elevation cold water habitat, including lakes, glacial-fed streams, and flowing springs in the Cascade Mountains (Kondratieff and Lechleitner 2002, Baumann and Kondratieff 2010, Kubo et al. 2013, Fallon and Blevins 2020). It should be noted that Lednian species may be particularly at risk from climate change impacts such as receding glaciers, reduced snowfall, loss of perennial snowpack, and warmer summer water temperatures (Giersch et al. 2016). For example, one species of *Lednia* (*L. tumana*) was recently listed as threatened under the Endangered Species Act as a result of glacial habitat loss under climate change (Muhlfeld et al. 2011, USFWS 2019).

Field collections of *Lednia* spp. have tended to occur only after years of surveys in remote, backcountry montane habitats. Yet despite their apparent rarity, members of the genus *Lednia* may be more widespread than currently known; limited observations may be due to difficulty of survey site access, low detection probability in part due to narrow adult activity windows, and public lack of interest in or knowledge of montane stoneflies. To determine if *L. borealis* was more broadly distributed in Washington than the scattered collections that had been previously reported suggest, additional surveys were necessary. We aimed to supplement existing information regarding the distribution, population trends, and conservation needs of this group of stoneflies. Therefore, the purpose of this study was to conduct targeted surveys within suitable habitat across the potential range of *Lednia borealis* in Washington State, including resampling of historic locations and documenting new populations. Additionally, detailed information on habitat, co-occurring aquatic invertebrate species, and timing of detection were collected to improve future survey and modeling efforts for documenting rare montane stoneflies in the Pacific Northwest.

## MATERIALS AND METHODS

### Species of interest

The northern forestfly *(Lednia borealis)* is a species of stonefly documented in Washington State, USA. Adults of the species are described as 5 to 7.5 mm long with dark brown body color and hyaline wings with darker veins near the cord (Baumann and Kondratieff 2010). Larvae can reach 4.5 to 6.5 mm and have tiny light-colored spines on their legs (Baumann and Kondratieff 2010). A distinguishing characteristic for *Lednia* larvae is the absence of gills, which can make identification of the genus somewhat easier in the field (Baumann and Kondratieff 2010, J. Giersch pers. comm. 2015). Past records for *L. borealis* indicate that adults are active from mid-July through mid-September, although one specimen was collected as early as June 1. Little is known about *L. borealis* feeding habits, although most species in the Nemouridae family are shredders or collector-gatherers and use a variety of coarse plant materials (DeWalt and Kondratieff 2019).

### Data review

Prior to selecting survey sites, we conducted a review of all published or digitally accessioned records for *Lednia* species in Washington, gathering data from GBIF, SCAN, Web of Science, Google Scholar, the Plecoptera Species File, GenBank, and BOLD Systems. We also requested species occurrence data from state and federal land management agencies in Washington, including US Forest Service Region 6, Oregon/Washington Bureau of Land Management, Washington Department of Fish and Wildlife, and the Washington Natural Heritage Program. Species experts and authors were consulted to ensure unpublished data were not missed in our review; these included Richard Baumann, Boris Kondratieff, Joe Giersch, and Scott Hotaling. Through our initial data review, we identified just 11 occurrence records at 9 sites for *Lednia borealis* spanning the years 1987 to 2014 in habitats ranging from springs to lakes and streams, and from 1,162 to 1,747 meters above sea level (Kondratieff and Lechleitner 2002, Baumann and Kondratieff 2010, J. Giersch pers. comm. 2015). However, following our surveys, an additional 2 occurrence records were identified in the Canadian National Collection (Table 1; Supplementary Materials).

**Table 1.** *Lednia borealis* collections referenced in Figure 1. Site code corresponds to Supplementary Material.

| Site Code | Main Waterbody | Collection Date | Collector | Collection |
| --- | --- | --- | --- | --- |
| CZ | North Fork Cascade River | 6/1/1965 | F. Schmid | 1 adult female |
| CY | Coleman Glacier | 8/12/1975 | J.M. & B.A.C | 1 adult female |
| CV | Upper Bagley | 7/30/1987 | M.M. Ellsbury | 1 adult female |
| CT | No Name Lake | 8/24/1989 | R. Hoffman | 3 adult female |
| CM | Redoubt Lake | 8/29/1989 | E. Deimling | 1 adult male |
| CU | Price Lake | 9/6/1989 | E. Deimling | 1 adult male |
| CJ | Middle Tapto Lake | 9/8/1989 | G. Lomnický | 1 adult male |
| CI | Nert Lake | 9/11/1989 | R. Hoffman | 1 adult female |
| CS | Fryingpan Creek | 7/16/2000 | B.C. Kondratieff and/or R.A. Lechleitner | 1 adult male |
| CG | Snow Lake spring | 2010: 8/23; 8/25; 9/8 | J. Kubo | 5 adult male; 9 adult female; 9 immature |
| C | Colchuck Lake | 8/8/2014 | S. Hotaling | none (visual) |
| BM | Lyman Lake | 8/11/2018 | E. Blevins | 1 adult female |
| BU | White Chuck River | 8/18/2018 | D. Stinson, T. Cotten | 1 adult female |
| CA | Green Lake | 8/28/2019 | C. Fallon, E. Blevins | 1 adult female |
| CB | Sholes Glacier | 8/28/2019 | C. Fallon, E. Blevins | 1 adult male; 2 adult female |
| CH | Unnamed tributary to Green Lake | 8/28/2019 | C. Fallon, E. Blevins | 3 immature |
| AL | Coleman Glacier | 8/29/2019 | C. Fallon, E. Blevins | 1 adult female |
| BX | Coleman Glacier | 8/29/2019 | C. Fallon, E. Blevins | 2 adult female |
| CC | Coleman Glacier | 8/29/2019 | C. Fallon, E. Blevins | 1 adult female; 3 immature |

**Figure 1.**
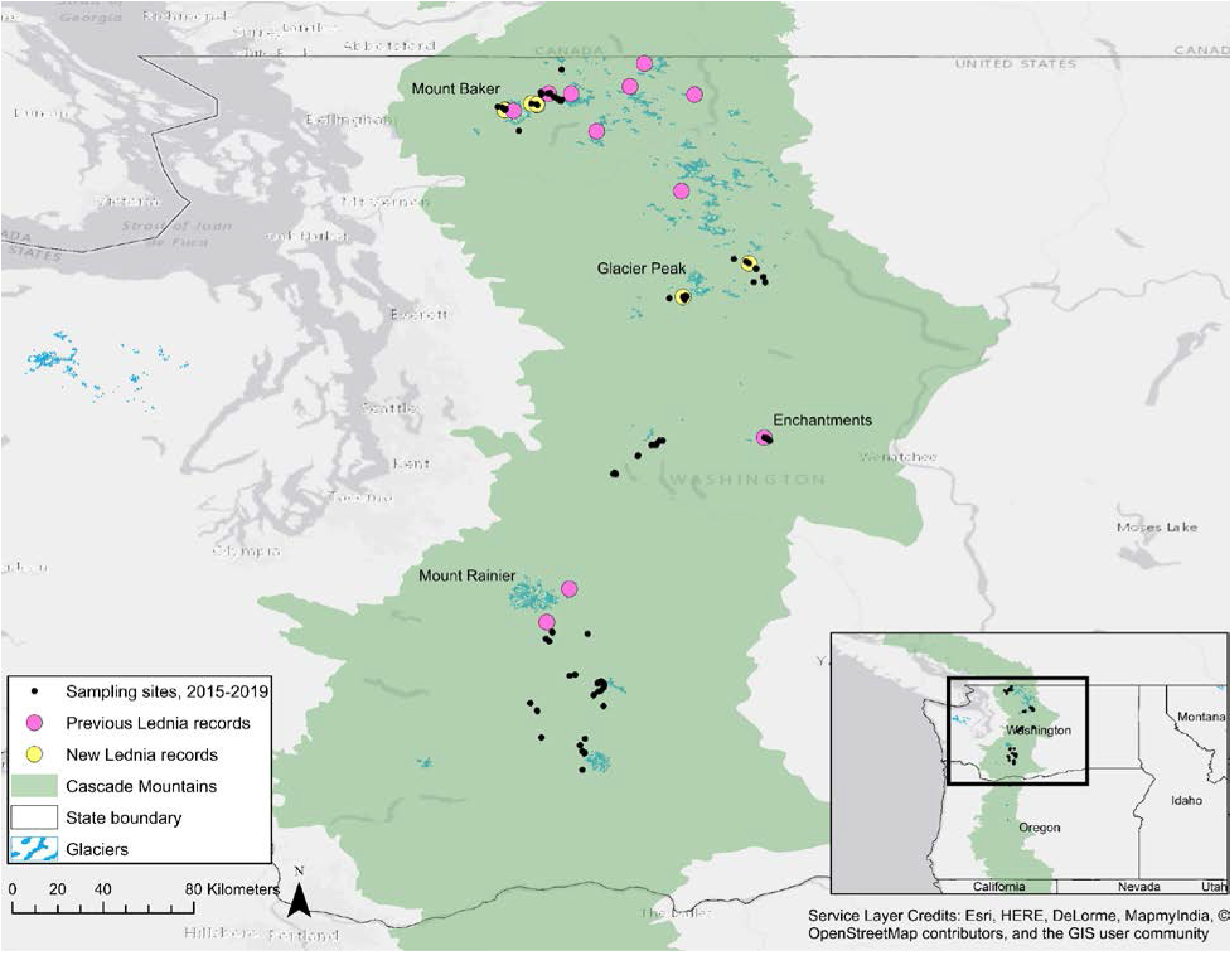
Locations sampled for *Lednia borealis* between 2015 and 2019 in the Cascade Mountain Range of Washington and all known *L. borealis* sites in the state.

### Site selection

Potential habitat on US Forest Service lands was first determined by mapping these known locations of *Lednia borealis* in ESRI ArcMap 10.4. Forest Service land ownership boundaries were overlaid with a National Hydrography Dataset (NHD) layer and a US Glacier Database (PSU 2005) layer for Washington State. All four layers (known locations, land ownership, hydrography dataset, and glacier layer) were converted to KML files and imported to Google Earth, which was used to select potential sites for surveys. The time-lapse Historical Imagery feature on Google Earth was used to determine which areas were most likely to be influenced by year-round snow (Giersch et al. 2016). All sites selected were on the Gifford Pinchot, Okanogan-Wenatchee, and Mount Baker-Snoqualmie National Forests. To the extent possible using aerial imagery and shapefiles, we prioritized alpine and subalpine areas with glacial outlets, small streams, seeps, and springs, while also considering relative ease of access via established hiking trails.

### Survey period

Surveys were conducted in July and August to help minimize exposure to potentially dangerous early-season stream crossings in the mountains and deep snow that persisted late into the season in the glacial valleys and north facing slopes. The survey window was within the majority of previous collection dates of *Lednia borealis*, which span mid-July through early September.

### Sampling methodology

All sites visited between 2015 and 2019 were surveyed by the authors or volunteers using standard recommended survey techniques for stoneflies. Stream, lake, and seep habitats were surveyed by overturning rocks, looking through the substrate, and sweep-netting and/or beating adjacent vegetation (when present). Surveyors spent a minimum of 20 person-minutes searching each site, which varied in size. The waterbody name, locality information, water temperature, substrate type(s), water depth, stream width, canopy cover, streamside vegetation, and time spent surveying were recorded at each site. Canopy cover was binned into three categories (exposed, exposed/forb, or vegetated canopy [e.g. forested]), representing distinctly different levels of exposure among sites. Site coordinates were marked using a Garmin Rino 120 or eTrex 20x unit (NAD83) in years 2015 to 2018. In 2019, surveyors used the S1 Mobile Mapper application on an Android phone, a custom mapping and field data collection tool built by the Service First (S1) Mobile GIS team (sponsored by the OR/WA Bureau of Land Management and USFS Region 6). Immature specimens were collected with forceps and placed in plastic screw-cap vials containing 95% ethanol as a preservative. Adults were collected from sweep nets or with forceps and preserved the same manner. Although stoneflies were the target, surveyors also collected caddisflies and occasionally mayflies. Surveys took place between 8:30 AM and 8:30 PM. Incidental sites (i.e. sites that looked interesting but were not the focal target of our surveys) were sampled opportunistically in a less robust way, with the typical data recorded for these sites including date, location, and water temperature.

Sampling for *Lednia borealis* was conducted in Washington State over five years (2015 to 2019). Using the information drawn from our data review and from each subsequent year of sampling, we refined our study areas to incorporate areas of greater likelihood for detection, including lower and higher elevation springs and streams, and particularly targeting inlets to alpine lakes, glacial meltwater edges, and the surface of glaciers or snowfields (high elevation sites). The goal in sampling less likely sites at lower elevations was to determine the distributional boundaries and habitat characteristics of stoneflies like *Lednia* as compared to co-occurring species.

### Taxa identification

Specimen collections were identified by aquatic macroinvertebrate taxa experts Robert Wisseman (Aquatic Biology Associates, Inc.), Jon Lee (Jon Lee Consulting, CA), and Richard Baumann (Brigham Young University). In addition to morphological IDs, samples from a subset of the stonefly and caddisfly specimens were sent to LifeScanner (http://lifescanner.net/) for COI barcoding, a technology that allows for DNA-based species identification. A male-female pair of *Lednia borealis* from Sholes Glacier was deposited at Brigham Young University. All remaining specimens are housed at the Xerces Society for Invertebrate Conservation in Portland, OR.

## RESULTS

### Survey effort and site characteristics

Between 2015 and 2019, sampling was conducted at 94 sites by the authors or volunteers, including lakes and associated water sources (n = 39), other streams (n = 45), seeps or springs (n = 6), and glaciers (n = 4) (Figure 1; Table 1; Supplementary Material). Active survey effort averaged 56 person-minutes per site, although active survey effort generally increased whenever stoneflies of any species were detected at a site. In an effort to determine the range of elevations over which the species might occur, both lower elevation (484 meters) and higher (2,376 meters) sites were visited, although all visited sites occur above the previously reported lowest elevation collection for the species. Average recorded water temperature for all sites was 10.2° C (max = 17.5° C), whereas at sites where *Lednia* was collected, temperatures ranged from 2.5 to 12.5° C. Sampling sites also varied in the extent of adjacent or overhead vegetative cover, which we categorized from lower (“exposed”) to higher (“low vegetation” and “forested”) to account for differences in light and wind exposure at sites (Figure 2). Different waterbody types were distributed across the spectrum, except that springs and seeps tended to occur at higher elevations, and glaciers occurred only at higher elevations.

**Figure 2.**
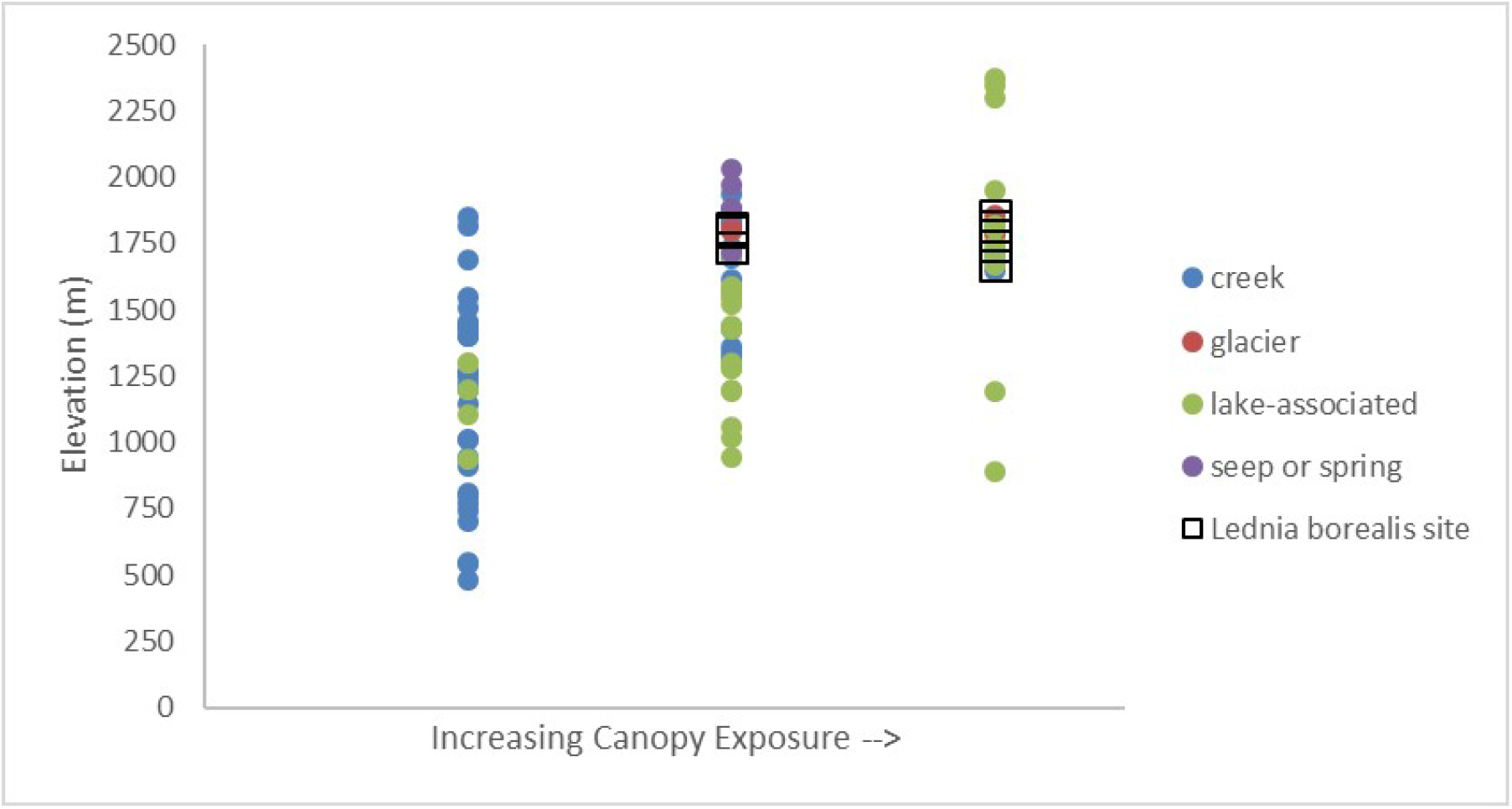
Characteristics of sampling sites (n = 94). Canopy exposure increases from left (“forested”), middle (“low vegetation”), to right (“exposed”).

### *Lednia borealis* collections

*L. borealis* was detected at 8 sites (3 of which are associated with a single glacier) during the study, bringing the total number of known locations to 19 (Figure 1; Supplementary Material). In this study, the species was detected only at creek (associated with snowmelt), glacier, or lake-associated sites, and never seeps or springs, despite prior collections from this habitat type by Baumann and Kondratieff (2010) (Figure 2). The species was also only detected at high elevation sites, occurring above 1,600 meters. At least one historic site (Upper Bagley Lake) was surveyed in three consecutive years but resulted in no detection of *L. borealis*, despite the species being detected elsewhere in the same week during 2019 surveys, but at elevations approximately 400 meters higher). Another recently reported site, Colchuck Lake, was visited during this study, although no *L. borealis* were detected. Although surveys were conducted in July, August, or September each year of the study, the species was only detected in August of 2018 and 2019. Collections included adult males and females as well as larvae; from these collections, six specimens of *L. borealis* were submitted for extraction and sequencing using LifeScanner kits. The results were then published to the Barcode of Life Data System (BOLD) (Ratnasingham and Hebert 2007). BOLD’s barcode index number (BIN) algorithm grouped all six sequences into the BIN BOLD:ACY8420, confirming that all six were indeed *L. borealis*. Additional aquatic invertebrates (primarily stoneflies and caddisflies, with a few mayflies) were collected at *L. borealis* sites, including representatives of 10 families and 12 genera (Supplementary Material). Although there was interest in comparing trends among co-occurring species, collections of the target species were too limited to draw any conclusions regarding co-occurrence of other caddisfly or stonefly species with *L. borealis*.

## DISCUSSION

Originally identified as *L. tumana* (a species found in Montana, USA, and Alberta, Canada), *L. borealis* is a relatively recently described species from Washington state, USA (Baumann and Kondratieff 2010, Giersch et al. 2016). Prior to the surveys described in this paper, *L. borealis* was reported from 11 sites in Chelan, Lewis, Pierce, Skagit, and Whatcom Counties, Washington (Kondratieff and Lechleitner 2002, Baumann and Kondratieff 2010, J. Giersch pers. comm. 2015, S. Hotaling pers. comm. 2019, CNC 2020), having first been collected in Washington state from Mineral Park (Skagit County) in Mt. Baker National Forest in 1965 (CNC 2020).

This study expands on the work conducted by Baumann and Kondratieff (2010), providing new insights into the current distribution of *L. borealis* in Washington, including new detections in the central (near Glacier Peak) and northern (near Mt. Baker) portions of the species’ known range, as well as a previously unpublished detection in the Enchantments (by S. Hotaling; J. Giersch pers. comm. 2015). It extends the upper elevational range from which this species is now known (highest site of detection = 1,860 meters), and reaffirms that habitat associations include glacial-edge habitat from Mt. Baker (Coleman and Sholes Glaciers).

Findings from this study suggest that *L. borealis* is more widespread than previously documented, but still quite rare. Specific habitat associations remain unclear, as the species was documented only at a subset of sites with representative habitat characteristics, such as availability of cold water, type of aquatic habitat, degree of shading or associated vegetation, elevation, and proximity to glaciers or permanent snowpack. Washington State is the most glaciated of the lower 48 United States, and the Glacier Peak Wilderness, which lies within this ecoregion, has more active glaciers than any other place in the lower 48 states (USFS 2016). As such, it is perhaps unsurprising that increased survey effort in the region has resulted in additional observations of the species, given the importance of glacier habitat to other species of *Lednia*, like *L. tumana* (Muhlfeld et al. 2011, Giersch et al. 2016). Yet the distance of *L. borealis* observations from existing glaciers appears to be highly variable among sites, from glacier edges up to several kilometers away, indicating that while the species may be affiliated with such habitat, it is not necessarily directly dependent upon it. It is possible that other alpine features, such as rock glaciers and subterranean ice, could play a role in this species’ distribution; increasing evidence shows that these features are highly important to mountain stream biodiversity (Brighenti et al. 2020, Tronstad et al. 2020). Regardless, the most abundant habitat for *L. borealis* in Washington is likely in the North Cascades ecoregion. Although only two populations were documented in the Glacier Peak Wilderness in 2018, surveys were far from comprehensive, and this area may well prove to be a stronghold for *L. borealis* populations. Additional known and potential habitat also occurs in North Cascades National Park, Mount Baker-Snoqualmie National Forest, and other mountainous wilderness areas of the state, including Mount Rainier National Park, Goat Rocks Wilderness, Alpine Lakes Wilderness, Mt. Adams Wilderness, and Olympic National Park. However, this study also suggests that Mount Rainier may form the southern end of the distribution. Indeed, despite seven years of relatively intense stonefly collecting in Mt. Rainier National Park by Kondratieff and Lechleitner (2002), this species was represented by only one collection (identified as *L. tumana*), and surveys conducted near the southern flank of Mount Rainier outside the national park as part of this study did not result in any detections, nor did surveys further south or east in the Cascade Range. To the north, the vast mountainous areas of British Columbia hold promise for this species. Recent evidence suggests that *Lednia borealis* occurs in the mountains of central British Columbia (Ratnasingham and Hebert 2013, Green et al. in preparation), representing a substantial range extension.

This study also highlights additional challenges to understanding the species’ current distribution. For example, surveys at two other previously-documented sites, one historic (Upper Bagley Lake) and one recent (Colchuck Lake), resulted in no detections. Additional surveys are needed to revisit historic sites to determine the status of populations, as it is unclear whether historic sites are still occupied, whether the phenology of adult emergence is changing, or whether sampling approaches limit detection of the species where it occurs. Indeed, few individuals were observed when the species was detected (i.e., fewer than 2 adults on average among sites visited in this study), sites are often remote and difficult to access, and the species may be active in its adult form (and thus more easily detected and identified) for only a week or two each year. It is recommended that surveys continue at historic, new, and potential sites to continue to improve understanding of the species and its alpine habitat, and future survey efforts that occur over longer periods of time in areas of glacial runoff may prove the most successful. Development of improved survey techniques also has the potential to help researchers understand and predict distribution of the species. As per the recommendations by Birrell et al. (2020), future surveys should also focus on collecting more abiotic information on high elevation sites.

Although relatively little information is available about this and other cold water-dependent stonefly species, what is known about better studied species, including *Lednia tumana* and *Zapada glacier*, shows cause for concern under the changing climate of alpine areas in western North America; both *L. tumana* and *Z. glacier* have recently been listed as threatened species under the Endangered Species Act (USFWS 2019), due primarily to heightened extinction risk because of climate change (Muhlfeld et al. 2011, Giersch et al. 2015, Giersch et al. 2016). Climate change in the region is causing glaciers to shrink and disappear and is altering snowpack trends and phenology (Pelto 2008), and although high elevation streams fed by snow and springs may continue to serve as critical climate refugia for cold water-dependent species even after glaciers disappear (e.g., Muhlfeld et al. 2020), without better understanding of this species’ habitat and needs in the alpine environment, it is unclear what future impact climate change may have on this and other alpine aquatic insects. As a result, additional surveys and study of *L. borealis* are critical for understanding its conservation needs.

## ACKNOWLEDGEMENTS

We acknowledge the Interagency Special Status/Sensitive Species Program for funding initial surveys by the Xerces Society for Invertebrate Conservation on Forest Service lands in 2015 and 2016. Survey data were provided by volunteer Todd Folsom in 2017, 2018, and 2019. Survey efforts in 2018 and 2019 were funded by the Northwest Ecological Research Institute. We thank Sarina Jepsen, Todd Folsom, David Green, and the many researchers who have provided records, collecting advice, and identification services over the years, including Jon Lee, Bob Wisseman, Scott Hotaling, Joe Giersch, Bill Stark, Boris Kondratieff, and Richard Baumann, and David Burton. We thank Chris Sato and the Washington Department of Fish and Wildlife for their support. We also thank Richard Vacirca of the U.S. Forest Service for permit assistance. Earlier drafts of this manuscript were improved by comments from Sarina Jepsen and Scott Hotaling.

## Supplementary Material

Table 1. Sites visited and their characteristics.

Table 2. All prior known Washington records for *L. borealis*.

Table 3. All species collected and identified in this study. Note that target species were stoneflies and caddisflies, all others incidental.

Figures. Photos of *Lednia borealis* collection sites, 2018 and 2019.

