## Supplementary Material - Figures for "New distributional data for the northern forestfly, *Lednia borealis* Baumann and Kondratieff, 2010 (Plecoptera: Nemouridae), Washington, USA"

**Site photos for 8 newly documented *Lednia borealis* sites in Washington State, USA, 2018-2019.** Site codes are provided in parentheses after each site name, and correspond to Supplementary Tables.

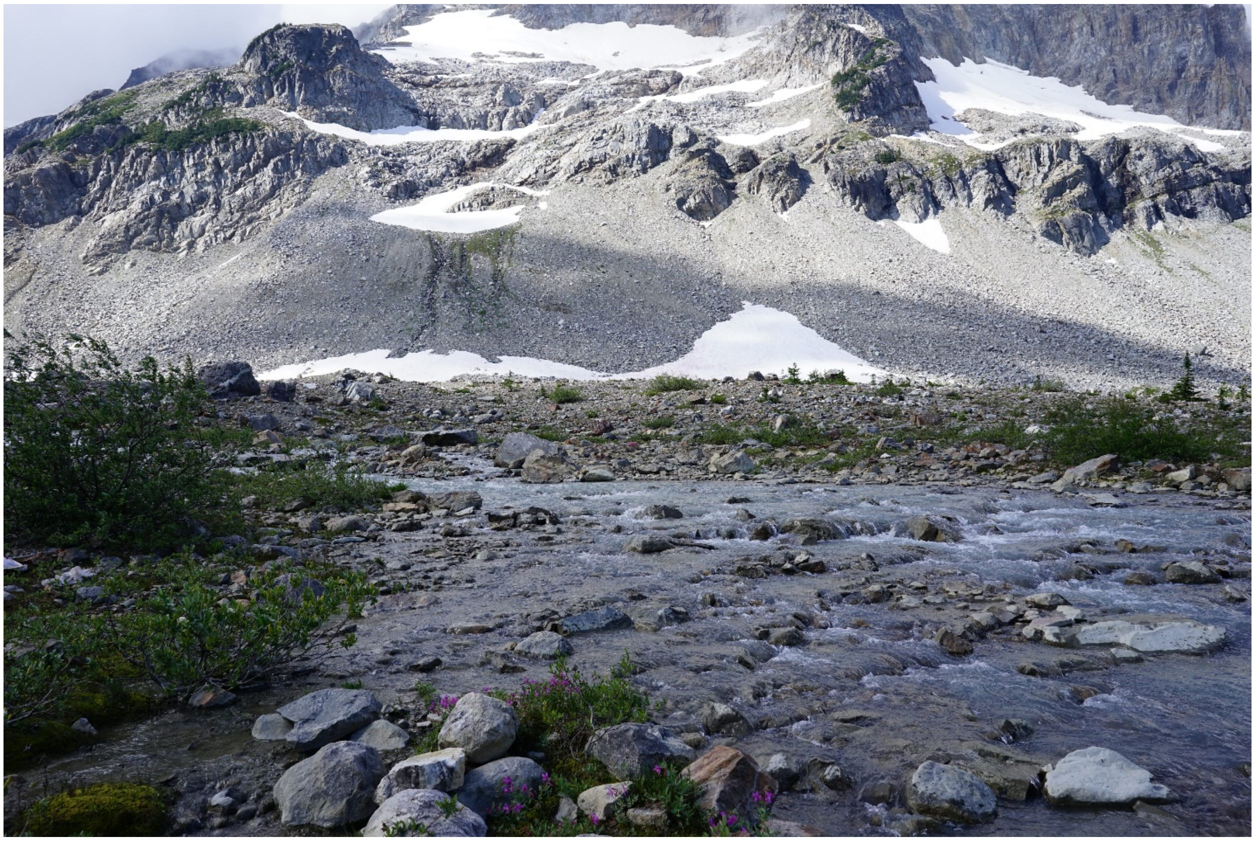

Upper Lyman Lake #2 (BM): August 11, 2018. Okanogan-Wenatchee National Forest, WA.

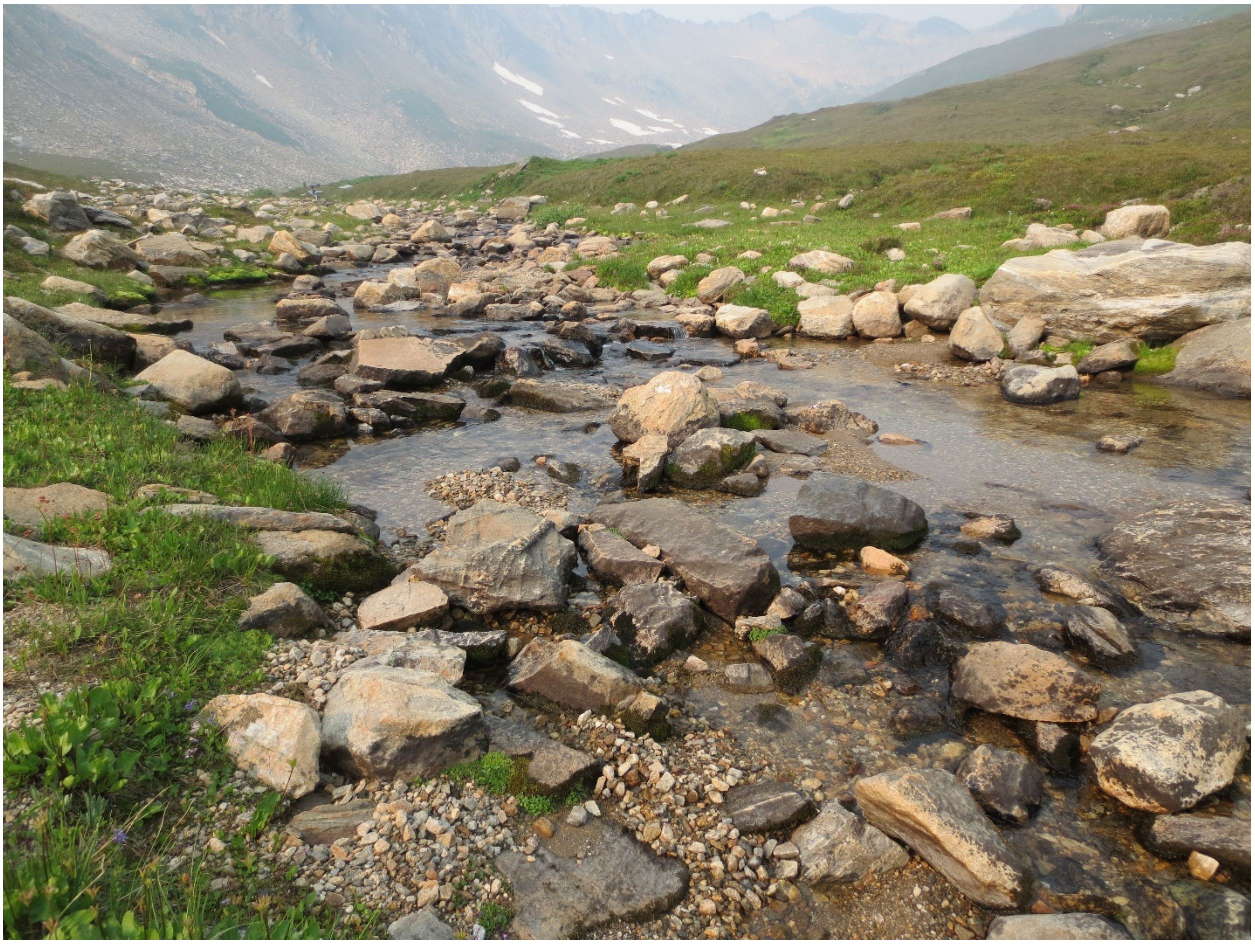

White Chuck Red Pass Fork (BU): August 18, 2018. Mount Baker-Snoqualmie National Forest, WA.

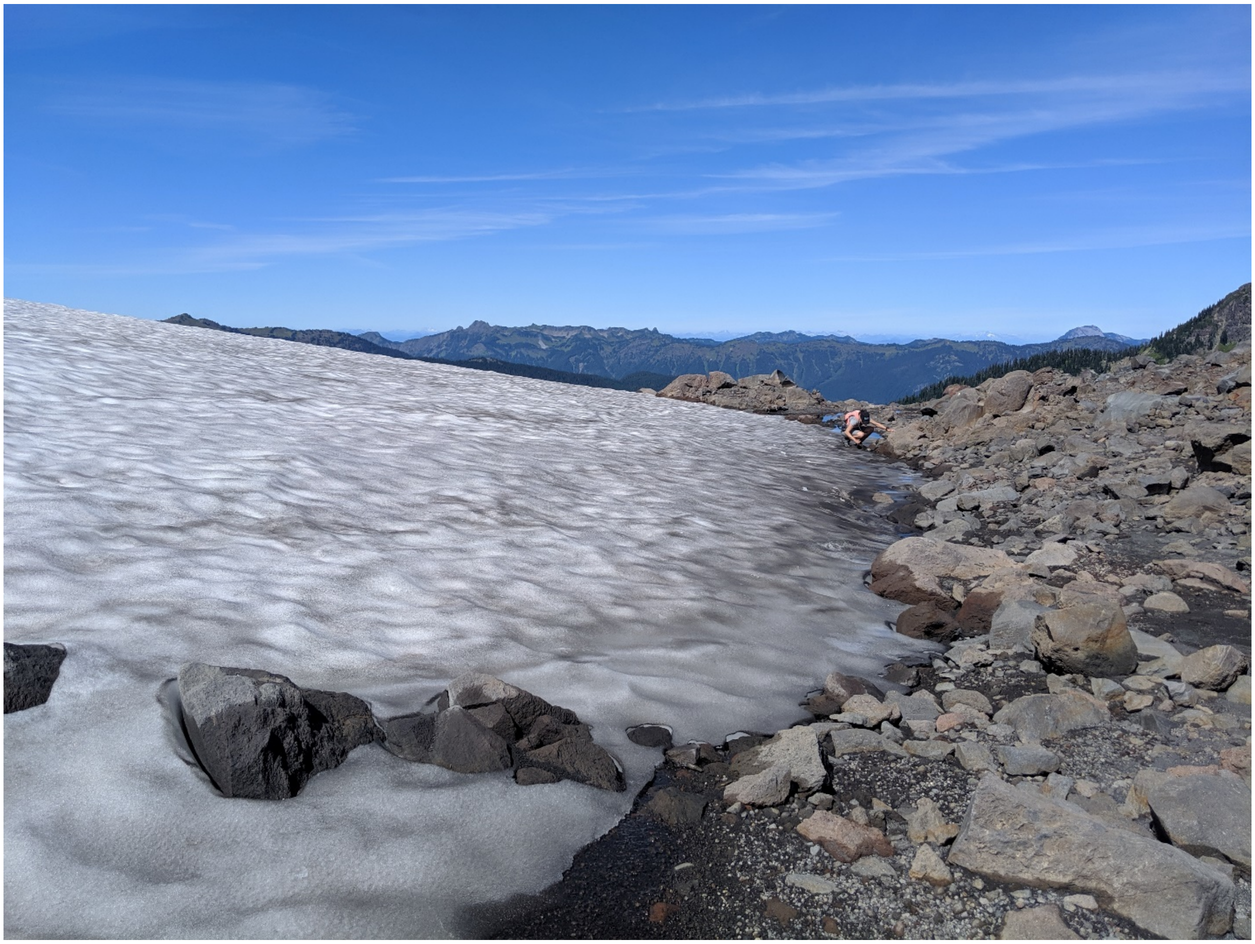

Sholes Glacier (CB): August 28, 2019. Mount Baker-Snoqualmie National Forest, WA.

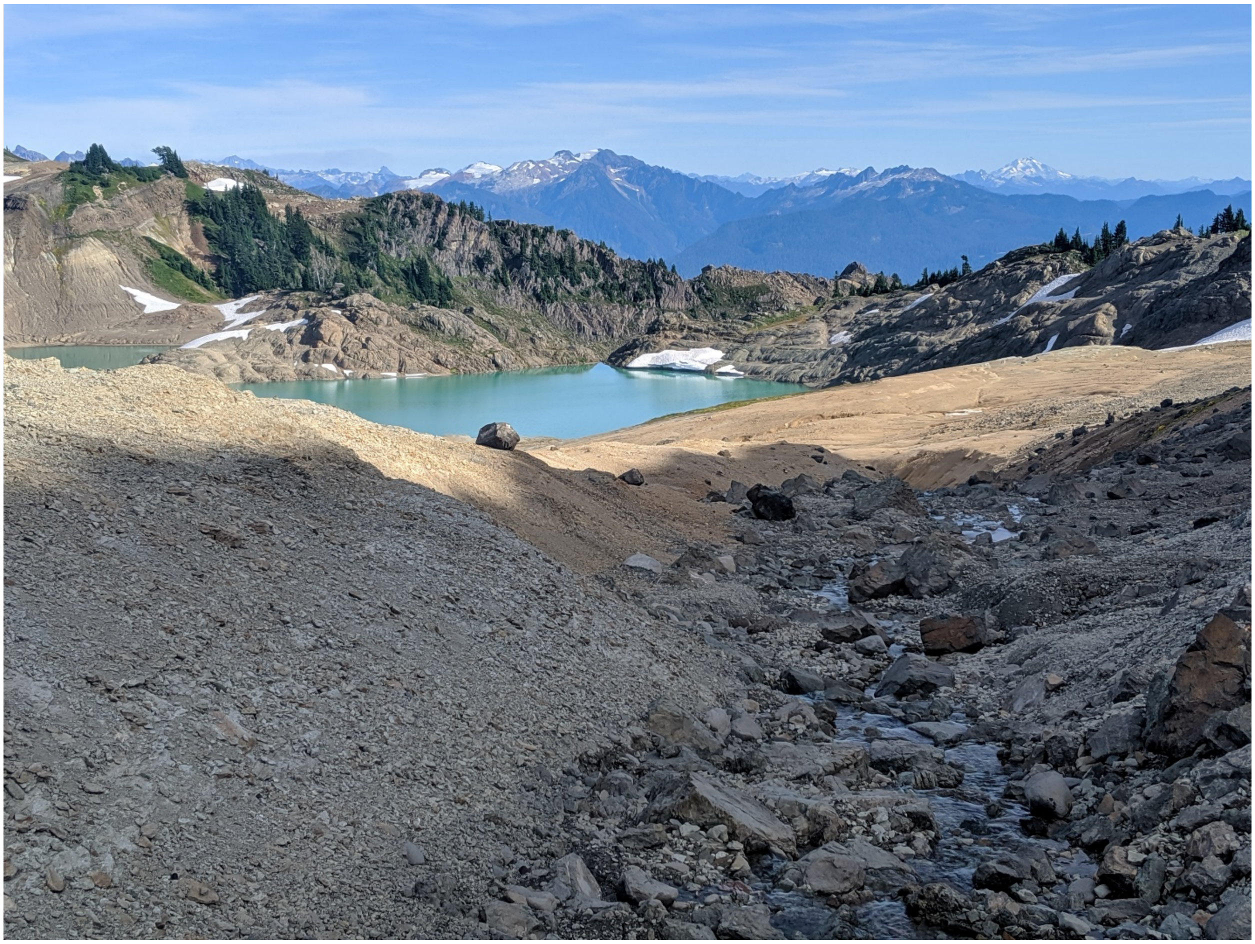

Green Lake inlet creek (CA): August 28, 2019. Mount Baker-Snoqualmie National Forest, WA.

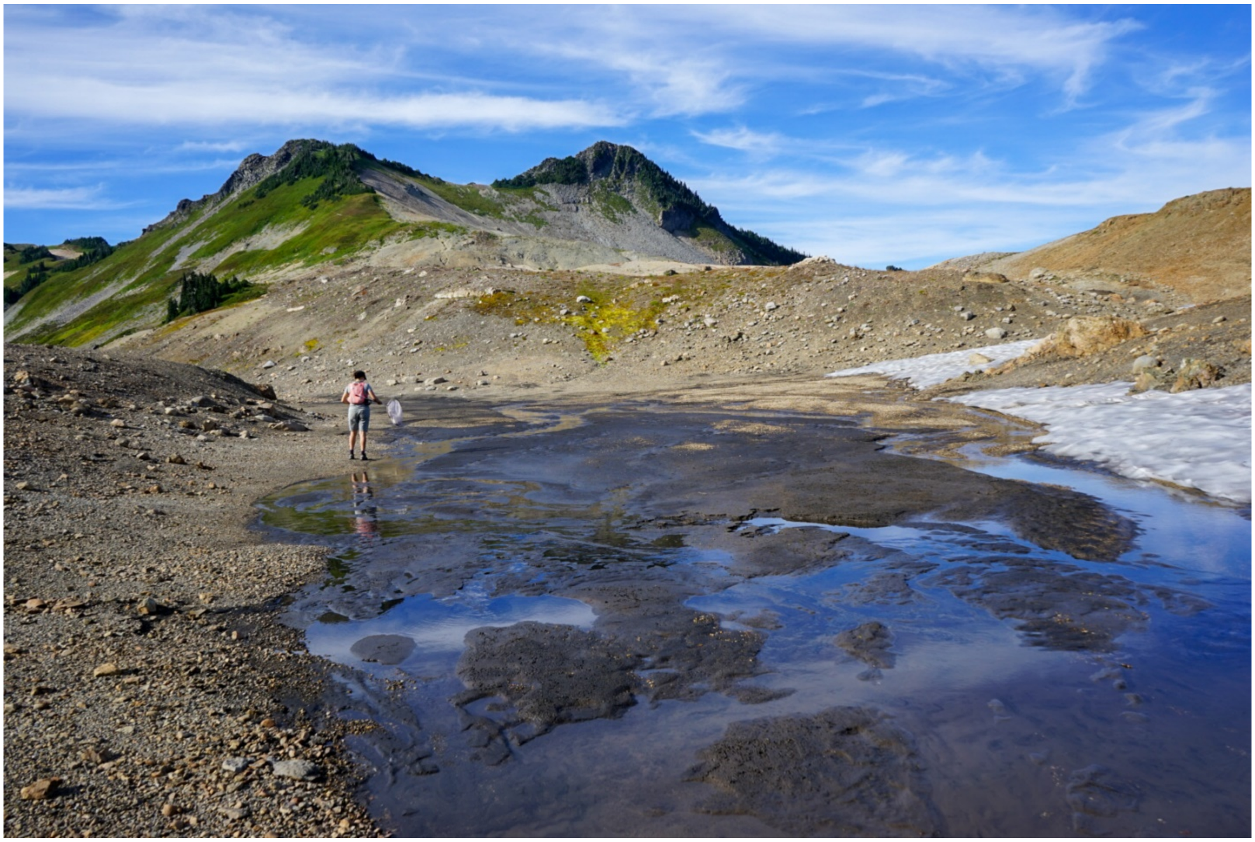

Unnamed tributary to Green Lake (CH): August 28, 2019. Mount Baker-Snoqualmie National Forest, WA.

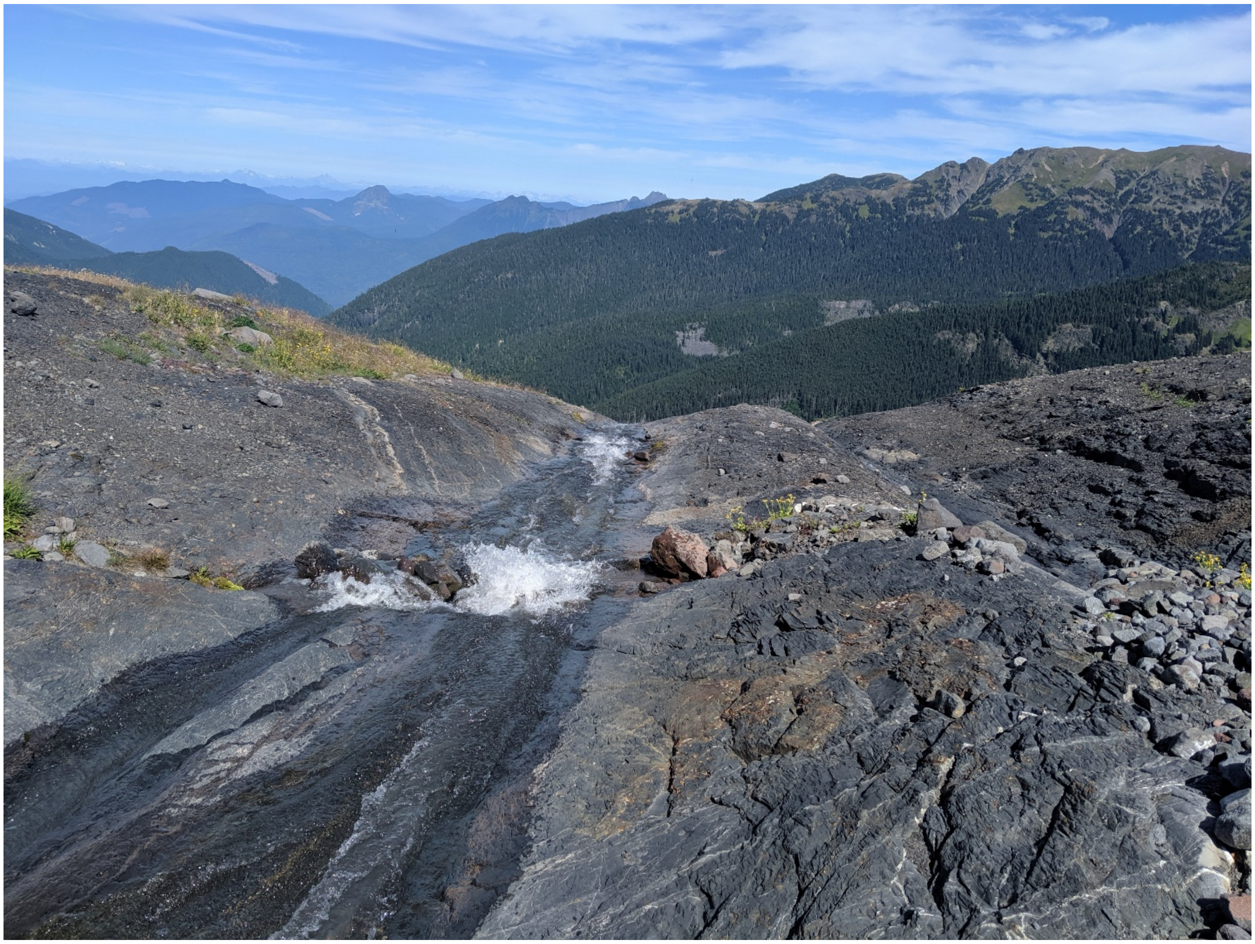

Coleman Glacier #1 (AL): August 29, 2019. Mount Baker Snoqualmie National Forest, WA.

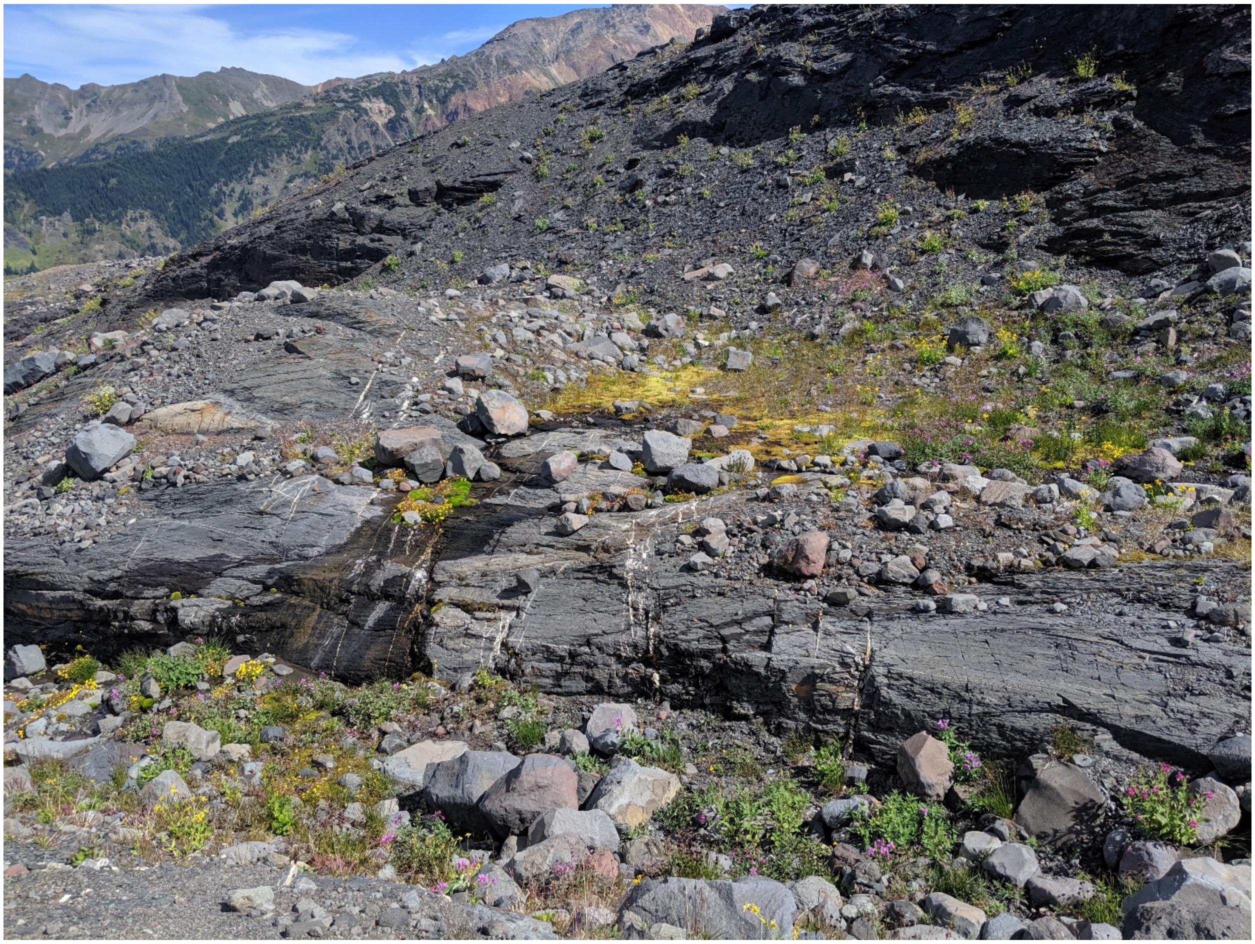

Coleman Glacier #2 (BX): August 29, 2019. Mount Baker-Snoqualmie National Forest, WA.

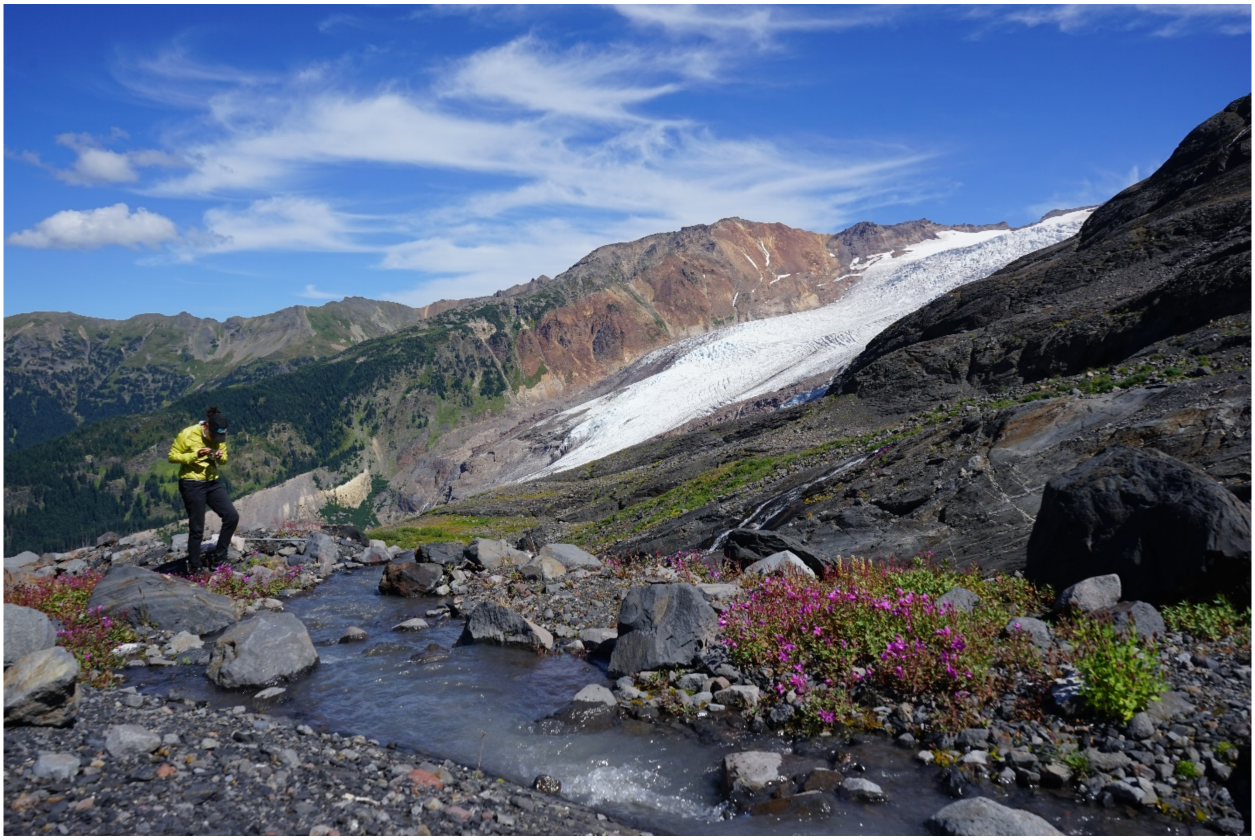

Coleman Glacier #3 (CC): August 29, 2019. Mount Baker-Snoqualmie National Forest, WA.
